# Evolution of phenotypic plasticity during environmental fluctuations

**DOI:** 10.1101/2023.01.22.523389

**Authors:** Zuzana Sekajova, Erlend I. F. Fossen, Elena Rosa, Irja I. Ratikainen, Manon Tourniaire-Blum, Elisabeth Bolund, Martin I. Lind

## Abstract

Evolution in fluctuating environments is predicted to disfavor specialization and instead select for alternative strategies, such as phenotypic plasticity or possibly bet-hedging, depending on the accuracy of environmental cues and type of fluctuations. While these two alternatives are often contrasted in theoretical studies, their evolution are seldom studied together in empirical work.

We used experimental evolution in the nematode worm *Caenorhabditis remanei* to simultaneously study the evolution of plasticity and bet-hedging in environments differing only in their temperature variability. We exposed worms for 30 generations to either fluctuating or slowly increasing temperature, these two environments had the same average temperature over evolutionary time. After experimental evolution, we scored size at sexual maturity and fitness in full siblings reared in two different temperatures, optimal 20°C and mildly stressful 25°C.

Experimental evolution in the fluctuating environment resulted in the evolution of increased body size plasticity but not increased bet-hedging, compared to evolution in the slowly changing environment. Plasticity followed the temperature size rule as size decreased with increasing temperature and this plastic response was adaptive. In addition, we documented substantial standing genetic variation in body size, which represents a potential for further evolutionary change.

## Introduction

Natural environments are generally not stable, but fluctuate on both long and short timescales, and a developing organism needs to take this environmental variation into account. While the parental environment can be a reliable cue if the environmental fluctuations are slow and predictable (Lachmann & Jablonka, 1996; Kuijper & Hoyle, 2015; Leimar & McNamara, 2015; Uller *et al*., 2015) and can result in the evolution of anticipatory parental effects (Dey *et al*., 2016; Lind *et al*., 2020), faster and random fluctuations select against inheritance of the parental phenotype. Instead, phenotypic plasticity or bet-hedging is predicted to be adaptive (Moran, 1992; Simons, 2011). If the developmental environment provides reliable cues for the optimal offspring phenotype, theory predict the evolution of adaptive phenotypic plasticity (Moran, 1992; Gavrilets & Scheiner, 1993; Simons, 2011) which is the ability of a genotype to produce different phenotypes depending on environmental conditions (West-Eberhard, 2003). If plasticity is present, individuals will canalize fitness between environments by adjusting their phenotype according to the environmental conditions.

Alternatively, individuals may express bet-hedging, which acts to reduce variation in fitness (especially to avoid very low fitness values in certain environmental states) at the cost of lowered arithmetic mean fitness, often by producing offspring with a range of phenotypes (diversified bet-hedging), where some of the offspring always matches the environment and is successful (Philippi & Seger, 1989). In contrast to phenotypic plasticity, bet-hedging is generally seen as favored when environmental cues have low predictability (Cohen, 1966; Slatkin, 1974; Meyers & Bull, 2002; Kussell & Leibler, 2005; Wolf *et al*., 2005), or when there is low correlation between the environment of development and selection (Tufto, 2015).

While both phenotypic plasticity and bet-hedging can be adaptive in variable environments (Simons, 2011; Furness *et al*., 2015), most empirical studies of evolution in variable environments focus on the evolution of plasticity. While phenotypic plasticity is common and well documented (DeWitt & Scheiner, 2004), very few studies have investigated the evolution of plasticity. These studies generally follow two lines, either they investigate whether increased plasticity evolves in more variable environments (Moran, 1992; Tufto, 2015), or whether increased plasticity evolves as a (possibly transient) step in adaptation to a novel but stable environment (Lande, 2009; Chevin *et al*., 2010; Levis & Pfennig, 2020). Studies focusing on the role of environmental heterogeneity have found stronger phenotypic plasticity in natural populations (Lind & Johansson, 2007; Lind *et al*., 2011) or species (Hollander, 2008) originating from more variable environments. Moreover, two recent experimental evolution studies in microalgae have shown that variable environments can select for increased phenotypic plasticity (Schaum *et al*., 2022), but that very unpredictable environments also can select against plasticity (Leung *et al*., 2020). One possible explanation for selection against plasticity during rapid fluctuations is that it may take time for the plastic phenotype to be induced (Burton *et al*., 2022; Dupont *et al*., 2024). Studies that instead focus on the evolution of plasticity during adaptation to new stable environments have found that plasticity can rapidly evolve in a new environment (Sikkink *et al*., 2014b; Corl *et al*., 2018), but also that maladaptive plasticity can play a major role (Ghalambor *et al*., 2015; Campbell-Staton *et al*., 2021).

In contrast to phenotypic plasticity, empirical studies documenting bet-hedging are rare (Simons, 2011), and the few examples suggest that it is more likely to evolve if the environments differ dramatically in fitness (Bull, 1987), such as delayed germination in desert plants (Philippi, 1993; Clauss & Venable, 2000; Venable, 2007), diapause in killifish (Furness *et al*., 2015) or experimental evolution in fluctuating environments with large fitness differences (Beaumont *et al*., 2009; Graham *et al*., 2014). However, bet-hedging has been suggested as a mechanism explaining the strain-specific variance in reproduction between natural isolates of the nematode *Caenorhabditis elegans* (Diaz & Viney, 2014).

One environmental factor that is known to result in evolutionary adaptations (Berteaux *et al*., 2004; Geerts *et al*., 2015), but also alternative strategies, is temperature. Not only is temperature gradually increasing due to the ongoing climate change (Berteaux *et al*., 2004), but climate change also results in increased temperature variability (Easterling *et al*., 2000; Palmer & Räisänen, 2002; Van Aalst, 2006) potentially favoring evolution of increased plasticity or possibly bet-hedging. Indeed, most documented responses to climate change in natural populations are caused by pre-existing plasticity, while genetic adaptation seems rare (Merilä & Hendry, 2014).

Among traits showing plastic responses to temperature, body size is of immense importance to reproductive fitness. For females, a large body generally translates into increased egg production, and also males often benefit from large size (Hedrick & Temeles, 1989; Andersson, 1994). Therefore, perhaps surprisingly, in warm environments organisms generally increase growth rate, accelerate maturation but mature at a smaller size, which is called the temperature-size rule (Ray, 1960; Atkinson, 1994; Verberk *et al*., 2021). This has sometimes been argued to be a passive by-product of other temperature-dependent processes (Atkinson, 1994; van der Have & de Jong, 1996; Forster *et al*., 2011). However, small size may actually be actively regulated and beneficial in warm environments (Fryxell *et al*., 2020; Verberk *et al*., 2021) for example by being advantageous for thermoregulation (Partridge & Coyne, 1997), or allowing better regulation of oxygen demand and supply ratio (Walczyńska *et al*., 2015; Verberk *et al*., 2021).

We set out to investigate whether exposure to fluctuating temperature results in evolution of increased phenotypic plasticity compared to evolution in an environment with slowly increased temperature and whether any plasticity in body size is adaptive, using experimental evolution in the nematode *Caenorhabditis remanei*. *C. remanei* has a fast generation time and, since it is dioecious, harbors substantial standing genetic variation which makes it ideal for experimental evolution studies (Teotónio *et al*., 2017). In addition, *C. remanei* has been shown to respond to manipulations in temperature (Sikkink *et al*., 2014a; Lind *et al*., 2020), has the ability to respond plastically to new environmental conditions (Lind *et al*., 2020), and the plasticity to withstand heat-shock can itself evolve (Sikkink *et al*., 2014b). Body size of *C. remanei* is under directional selection towards an optimum larger than the mean size under standard temperature conditions (Stångberg *et al*., 2020), and pharmacologically lowered body size results in lower female reproduction (Lind *et al*., 2016). It’s close relative *C. elegans* also follow the temperature-size rule (Kammenga *et al*., 2007). As an alternative to phenotypic plasticity, we also investigated if populations evolving in variable environments had evolved increased diversifying bet-hedging.

We used previously established experimental evolution populations of *C. remanei* (described in Lind *et al*., 2020). During experimental evolution, replicate populations were exposed to 30 generations in one of two regimes; *Fast temperature cycles* or *Increased warming*. The *Fast temperature cycles* regime were switched between two temperatures (20°C, 25°C) every second generation, and although the fluctuations are deterministic, it represents an uncorrelated (and therefore unpredictable) fluctuating environment each generation, as the next generation will either be in the same or in a different temperature. The evolution in this environment was compared to the *Increased warming* regime, where worms were exposed to experimental evolution in a gradually increasing temperature which slowly raised from 20°C to 25°C over 30 generations, and which served as a control. Importantly, these two regimes had the same average temperature (22.5°C) over evolutionary time, and only differed in the rate and predictability of environmental change. While no theoretical model to our knowledge have investigated evolution of plasticity in fluctuating versus slowly changing environments, it corresponds to different degrees of fluctuation, which is well explored theoretically (Moran, 1992; Gavrilets & Scheiner, 1993; Tufto, 2015). After the 30 generations of experimental evolution, we reared full-siblings in either standard 20°C, or warm 25°C, and scored them for reproduction and body size.

We predict that worms evolving in fluctuating temperature every 2^nd^ generation would evolve increased phenotypic plasticity (relative to the *Increased warming* regime), since the timescale of this environmental variation is well within the parameter space where plasticity (and bet-hedging) is favored (Tufto, 2015). We predict that evolution of phenotypic plasticity is more likely than the evolution of bet hedging, since the differences in fitness between the two temperatures is likely to be relatively small (Lind *et al*., 2020). If increased phenotypic plasticity has evolved, we predict that the *Fast temperature cycle* populations would show increased size difference between temperatures, but not increased phenotypic variance within one temperature. If instead increased diversifying bet-hedging had evolved, we predict that the *Fast temperature cycle* populations will show (1) increased within-family variance within each temperature, and (2) decreased heritability of size, because of an increased environmental component of phenotypic variance (Tufto, 2015). We also predict that any plasticity in body size will follow the temperature-size rule, and that this plasticity is adaptive.

## Methods

### Experimental evolution

For the experimental evolution we used *C. remanei* nematode worms, strain SP8 which has been lab adapted for 15 generations at 20°C and subsequently exposed to 30 generations of experimental evolution in two regimes (*Increased warming* and *Fast temperature cycles*). The experimental evolution has been previously described in detail in Lind *et al*., (2020b). Briefly, in the *Increased warming* experimental evolution regime, the temperature gradually raised from 20°C to 25°C, which is a novel and mildly stressful temperature. This gradual change over 30 generations represent an increase of 0.1°C every 2.13 days and results in a correlated parental and offspring environment. In the second regime, *Fast temperature cycles*, the temperature fluctuated every second generation between 20°C and 25°C, resulting in 14 temperature shifts but no exposure to the intermediate temperatures. The environmental change is deterministic (every second generation) but since parents and offspring would end up in either the same or in different temperature, it represents uncorrelated parental and offspring environment. The generation time in *C. remanei* is temperature dependent; 4 days long in 20°C and 3.4 days long in 25°C. Despite these differences, the average temperature and the total chronological time of experimental evolution were identical for both regimes, at 22.5°C and 110 days respectively.

Each evolutionary regime consisted of six replicate populations. The populations were maintained on 92 mm Petri dishes poured with NGM agar in climate chambers set to 60% relative humidity. In order to prevent bacterial and fungal contamination, the agar and bacterial LB contained the antibiotics streptomycin and kanamycin and the fungicide nystatin. The plates were seeded with 2 ml of an antibiotic-resistant OP50-1 (pUC4K) strain of *E. coli* (Stiernagle, 2006) that served as a source of food. Every 1-2 days, a piece of agar containing approximately 150 worms of mixed ages was cut and transferred to a new plate containing fresh bacteria. This resulted in populations with overlapping generations that were maintained in a constant exponential growth phase. After the experimental evolution, populations were expanded for two generations and frozen in -80°C for later revival and subsequent phenotypic assays.

### Experimental set-up

Each replicate population of each of the two selection regimes was run in a separate block resulting in 12 experimental blocks in total. For logistic reasons we focus on females, since they are responsible for population growth rate and their fitness is straightforward to measure.

Briefly, populations were revived from freezing and exposed to 25°C for 3 generations, to avoid any maternal effects associated with freezing. The third generation of worms were split into eight families, each family consisting of one male and one female worm. From each family, we randomly picked eight offspring females (full siblings) and placed four females in 20°C and four in 25°C. Since our focus was evolution in females, their fitness was assessed by mating them with standardized males from the ancestral line. Females were kept together with two males (in case one of the males would be infertile/escape from the plate). The ancestral line was, in contrast to selection regimes, maintained for three generations in 20°C and subsequently split into 20°C and 25°C together with studied females. For the detailed description of the experimental set up, see supplementary figure 1. In total, we used 48 families per evolutionary regimes, and for each family we set up 4 offspring in each temperature.

### Phenotypic assays

#### Daily reproduction

Female and male worms were transferred to a new plate every 24 hours. The old plate was kept for two days to allow eggs to hatch and afterwards the number of viable offspring was counted to determine age-specific reproduction and calculate individual fitness. In the case of the males dying or escaping from the plate, they were replaced by a new male of the same age from the ancestral line. The female worm was discarded after dying, or after three consecutive days of zero reproduction.

#### Body size

The body size of the worm changes with time, reaching a peak before it declines (Lind *et al*., 2016), and the age at maximum body size is temperature dependent. Worms in 20°C reach their peak size at day 4 of adulthood (Lind *et al*., 2016). The peak size in 25°C is on day 2 of adulthood, which was determined during pilot assays. Photographs of worms were taken during their peak size using a Lumenera Infinity2-5C digital microscope camera mounted on a Leica M165C stereomicroscope. Body size was measured from the photographs using *ImageJ 1.46r* (https://imagej.nih.gov/ij/) as total cross-section area.

### Statistical analyses

All statistical analyses were conducted in R 3.6.1 (R Core Team, 2015).

#### Individual fitness

We used the age-specific reproduction data to calculate rate-sensitive individual fitness λ_ind_ for each individual (Brommer *et al*., 2002), which is analogous to the intrinsic rate of population growth (Stearns, 1992). Individual fitness was calculated by constructing a population projection matrix for each individual, and then calculating the dominant eigenvalue of this matrix, following McGraw & Caswell, (1996). Since we kept the population size and age structure constant during experimental evolution, individual fitness is maximized during evolution and is therefore the most appropriate fitness measure for this study (Mylius & Diekmann, 1995).

#### Thermal reaction norms

To test whether the degree of phenotypic plasticity has evolved, we used linear mixed-effect models to separately estimate the thermal reaction norms of body size and individual fitness, using the package *lme4* (Bates *et al*., 2015) in R. The models included either body size (area) or individual fitness (λ_ind_) as response variables. The full model included three fixed effects: mean-standardized temperature as a covariate, the experimental evolution regime as a categorical factor, and an interaction between temperature and evolution regime. We expect this interaction to be significant if the degree of plasticity has evolved during experimental evolution. Experimental line and dam identity were included as random effects. Significance of the fixed effects was evaluated using Wald χ^2^ tests. Pseudo-R^2^ values were calculated as the squared correlation coefficient between fitted values from the model and observed values.

#### Selection

To test if temperature responses in size are adaptive, we estimated the selection on body size and compared it to the observed temperature response. Selection on body size (area) was estimated using mixed-effect models in R with the package *lme4* and individual fitness (λ_ind_) as the response variable. The full model included the following fixed effects: area, area^2^, temperature (as a categorical factor with 2 levels), experimental evolution regime and all interactions between these variables except for interactions involving both area and area^2^ together. Experimental line was included as random effect, assumed to only affect the variance of the intercept. Significance of fixed effects was evaluated using Wald χ^2^ tests. Pseudo-R^2^ values were calculated as the squared correlation coefficient between fitted values from the model and observed values. The optimal size (i.e. the area that maximizes fitness) was calculated as: -b/(2×c), where b = the slope of the regression (i.e. the linear selection gradient) and c = the squared term of the regression (i.e. the quadratic selection gradient). Confidence intervals of the temperature-specific optimal sizes were generated by bootstrapping, implemented in the *boot* package using 10 000 bootstrap replicates.

#### Within family coefficient of variance

To test whether the degree of diversifying bet-hedging has evolved, we tested whether the experimental evolution regimes differed in the mean within family variance within temperatures. For each family, we used the trait values of the offspring (within a temperature) to calculate the within family variance. To account for differences in the means of the traits, we used within family means (i.e. the mean trait value of the family’s offspring) to mean-standardize the variance by calculating the within family coefficient of variance (CV): CV = σ/μ, where σ = within family standard deviation, and μ = within family mean. An ANOVA was used within each temperature to test if the evolution regimes differed in their mean within family CV.

Since it is more difficult to detect differences in variances than differences in means, we also performed power calculations on our ability to detect whether within-family CVs differ between the selective regimes. Balanced one-way ANOVA power calculations were performed to estimate the effect sizes possible to detect with power ranging from 0.70 to 0.95. Effect sizes, η^2^, were obtained for our sample size of N = 48 per selection regime and a significance level of 0.05. η^2^ is calculated as the sum of squared explained by the treatment (here: selection regime) divided by the total sum of squares, and has an equivalent interpretation as an R^2^.

#### Genetic variance and correlations

For body size (cross-section area), genetic variance and genetic correlations across temperature were estimated using animal models using the package *MCMCglmm* (Hadfield, 2010) in R. Univariate models were used to estimate genetic variance, whereas bivariate models were used to estimate genetic correlations, both using Gaussian family for trait distribution. An inverse Wishart prior with parameters V = 1 and nu = 0.02 were used in both univariate and bivariate models. Pedigree data linking offspring to parents, based on full-sib relationships, was included in the models. Convergence of the models were ensured by evaluating diagnostic plots of posterior distributions, using the convergence diagnostic half-width test by Heidelberger and Welch (1983), and by ensuring that the autocorrelation between MCMC samples was close to zero.

For univariate models, body area was used as the response variable. Temperature (as a categorical factor with 2 levels), experimental evolution regime, and an interaction between the two, were included as fixed effects. Genetic variance (V_G_), variance due to differences between experimental lines, and residual variances were estimated separately as random effects in the full model for each temperature-by-evolution regime combination. The full model ran for 4.2×10^6^ MCMC iterations, 0.2×10^6^ samples were discarded as burnin, and the thinning interval was 4000, resulting in a sample size of 1000 MCMC-samples. Reduced models, subset by temperature-by-evolution regime combination, were used to assess the statistical significance of V_G_, by comparing the deviance information criterion (DIC) of models with versus without genetic variance included.

Broad sense heritability (H^2^ = V_G_/V_P_, where V_P_ = total phenotypic variance after accounting for variance due to experimental line effects) and broad sense evolvability (I^2^ = V_G_/mean^2^, Hansen *et al*., 2003, 2011) were used to estimate the population’s evolutionary potential of body size. This was estimated for each temperature-by-evolution regime combination. Evolvability measures the expected percentage change in a trait per generation under unit strength of selection. Compared to heritability, evolvability is independent from the environmental variance and represents a measure of the evolutionary potential that is comparable across traits, populations and species when applied to traits with a natural zero and which are strictly positive (Hansen *et al*., 2011).

Genetic correlations of body size were estimated using bivariate animal models in *MCMCglmm.* The data was split in two subsets based on the experimental evolution regimes, resulting in genetic correlations being estimated separately for the two evolution regimes. Body area was the response variable and was treated as two traits in the models, as area at 20°C and area at 25°C. Random effects in the full models included genetic covariance between the two temperatures, whereas V_G_, variance due to differences between experimental lines, and residual variances were estimated separately for each temperature. The full models ran for 2.05×10^6^ MCMC iterations, 0.05×10^6^ samples were discarded as burnin, and the thinning interval was 2000, resulting in a sample size of 1000 MCMC-samples. Reduced models without genetic covariance were used to access the statistical significance of the genetic covariance, by comparing the DIC of models with versus without genetic covariance included. The genetic correlation of body size across temperatures was calculated by dividing the genetic covariance by the product of the genetic standard deviation of the two temperatures. This was done on the posterior distributions, in order to carry the error forwards in the analyses.

To compare posterior distributions of H^2^, I^2^ and genetic correlations across temperatures and selection regimes, we calculated, within each MCMC sample, the pairwise differences in these measures and checked if the posterior distributions of these differences had a 95% credibility interval that included zero. Pairwise comparisons of distributions were only performed between evolution regimes within temperature, and between temperatures within evolution regimes.

## Results

### Thermal reaction norms

#### Size

Size decreased significantly with increasing temperature (Wald χ^2^ = 309.93, df = 1, p < 0.001, Fig. 1A). There was a significant interaction between temperature and evolution regime, where *Fast temperature cycles* had a steeper slope, meaning that it had evolved increased plasticity in size (Wald χ^2^ = 5.82, df = 1, p = 0.016). However, the intercepts (representing size at the mean temperature) were not significantly different between evolution regime (Wald χ^2^ = 0.09, df = 1, p = 0.769). The models pseudo R^2^ = 0.51. Variance components: V_dam_ = 25.03, V_Line_ = 20.92, and V_residual_ = 108.19.

**Figure 1.**
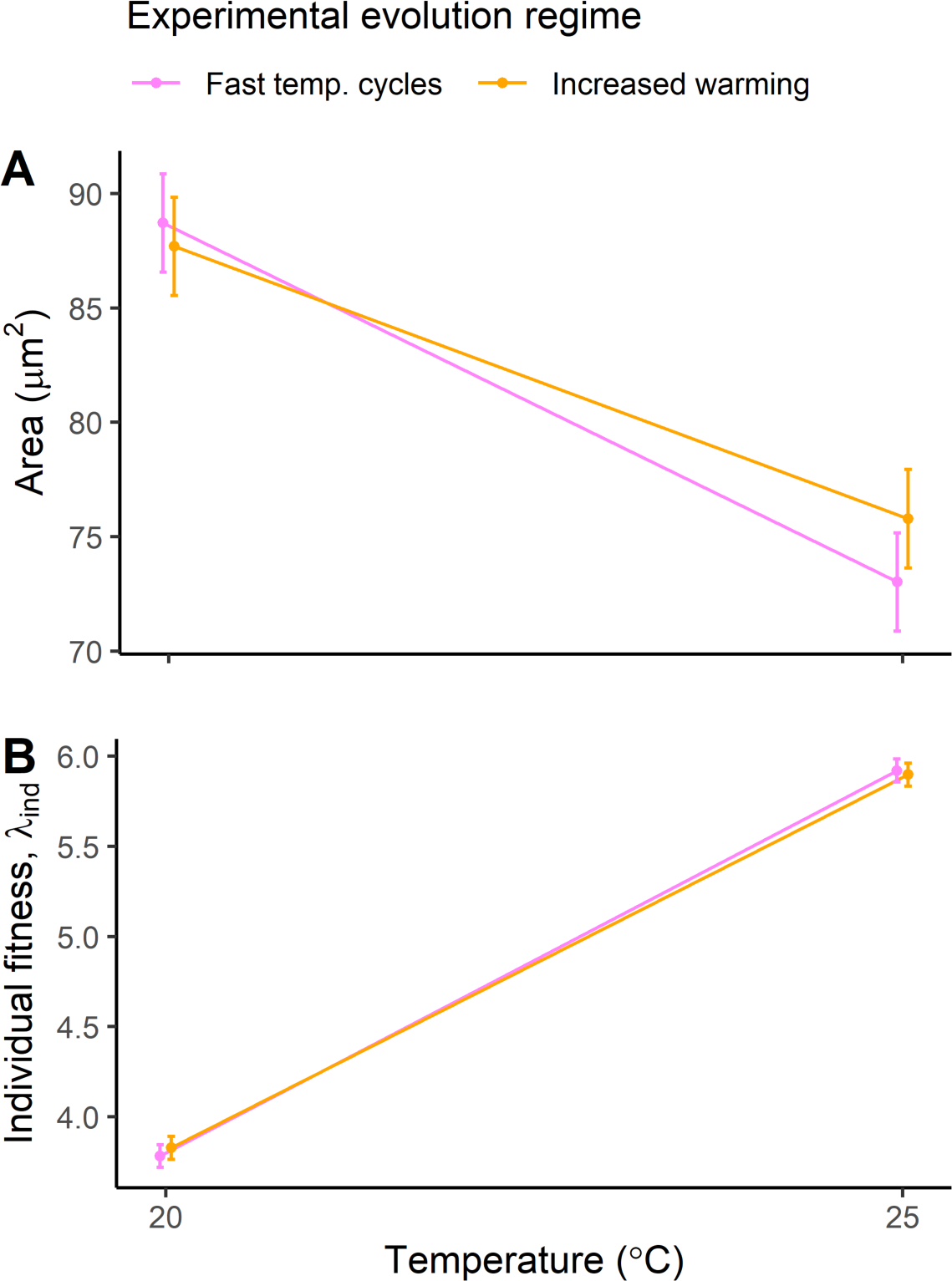
Thermal reaction norms showing means ± SE. (A) regression lines (with mean-standardized temperature): Fast; Area = 80.87 ± 2.08 – 3.14 ± 0.22 × Temperature. Increase; Area = 81.74 ± 2.08 – 2.38 ± 0.22 × Temperature. (B) regression lines (with mean-standardized temperature): Fast; λ_ind_ = 4.85 ± 0.06 + 0.43 ± 0.01 × Temperature. Increase; λ_ind_ = 4.86 ± 0.06 + 0.41 ± 0.01 × Temperature.

#### Individual fitness (λ_ind_)

The mean total reproduction decreased with temperature (mean ± SE: 20°C, 780 ± 17; 25°C, 672 ± 17; p < 0.001 for the difference between the temperatures). However, the individual fitness (λ_ind_) increased significantly with increasing temperature (Wald χ^2^ = 3670, df = 1, p < 0.001, Fig. 1B). The evolution regimes did not differ significantly in intercepts (Wald χ^2^ = 0.02, df = 1, p = 0.887), nor was there a significant interaction between temperature and evolution regime (Wald χ^2^ = 0.97, df = 1, p = 0.324). The best fit models pseudo R^2^ = 0.86. Variance components: V_dam_ = 0.05, V_Line_ = 0.01, and V_residual_ = 0.21.

#### Selection

There was significant linear and quadratic selection on body size (linear slope: Wald χ^2^ = 59.4, df = 1, p<0.001. Quadratic term: Wald χ^2^ = 40.3, df = 1, p < 0.001). The selection differed significantly between temperatures (Fig. 2), given by a significant overall temperature effect (Wald χ^2^ = 2992, df = 1, p < 0.001) and significant interaction effects between temperature and size (linear slope: Wald χ^2^ = 13.3, df = 1, p < 0.001. Quadratic term: Wald χ^2^ = 12.8, df = 1, p < 0.001). Maximum individual fitness (i.e. the optimal size) is predicted to be 93.73 µm at 20°C [95% bootstrap CI: 87.61, 112.98], and 84.19 µm at 25°C [95% bootstrap CI: 79.81, 92.61]. Selection was however not significantly different between the experimental evolution regimes (p > 0.22 for main effect and interactions between evolution regime and temperature or body size). No 3-way interaction between temperature, body size and evolution regime were significant (p > 0.18). The best fit model’s pseudo R^2^ = 0.85. Variance components: V_Line_ = 0.016, and V_residual_ = 0.204.

**Figure 2.**
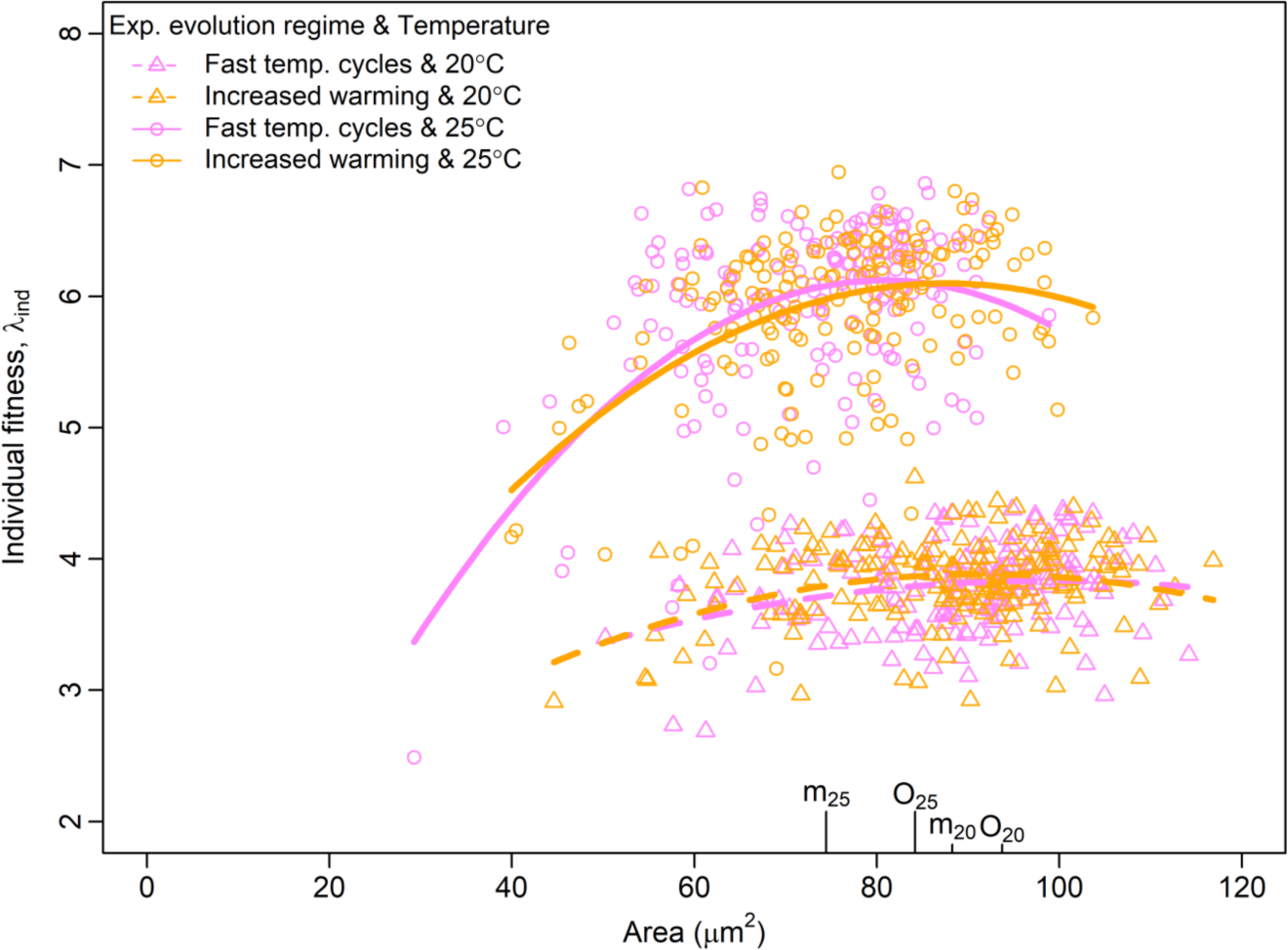
Selection on body size (area). Experimental evolution regimes were not statistically different, but are shown with separate lines. Overall regression line for 20°C: λ_ind_ = 1.596 ± 0.818 + 0.048 ± 0.020 × Area – 2.57×10^-4^ ± 1.16×10^-4^ × Area^2^. Overall regression line for 25°C: λ_ind_ = 0.138 ± 0.614 + 0.142 ± 0.017 × Area – 8.43×10^-4^ ± 1.20×10^-4^ × Area^2^. The mean size per temperature (m_20_, and m_25_) and optimal size (O_20_, and O_25_) are shown for 20°C and 25°C respectively. Individual fitness is higher in 25°C due to decreased development time, even if total reproduction is lower.

#### Within family CV

The evolution regimes did not differ significantly in within family CV of body size or of individual fitness at either temperature (Table 1). Moreover, the distributions of within family CV overlapped considerably between evolution regimes (Fig. 3). Power calculations showed that we had a power of 90% to detect effects where the evolutionary regimes explained at least 10% of the variation in within family CV (supplementary figure 2).

**Figure 3.**
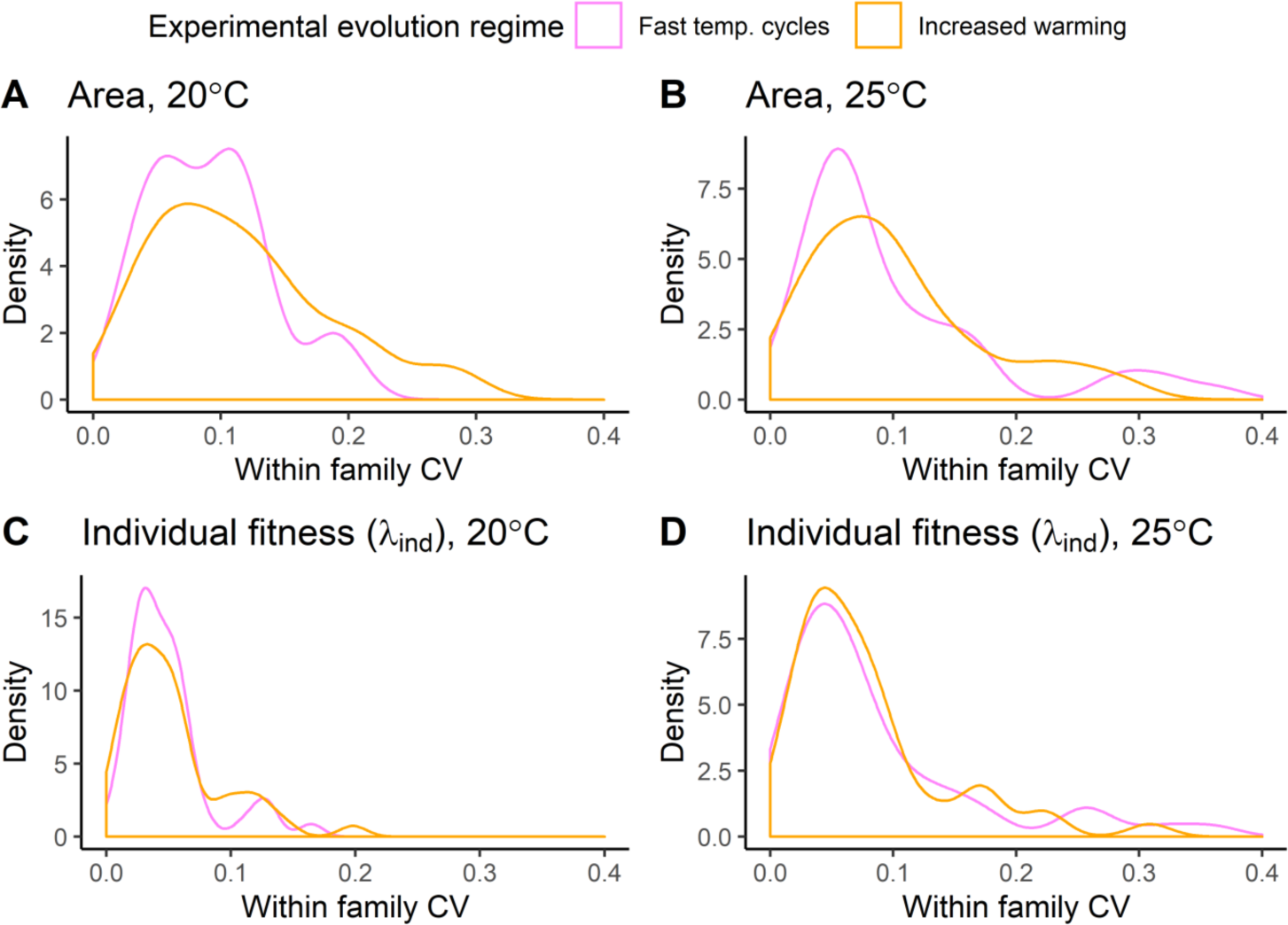
Distribution of within family coefficient of variance (CV) for two traits at two temperatures. The within family CV was estimated within temperature for each family by estimating the standard deviation (σ) and the mean (μ) of the family’s offspring trait values, where CV = σ/μ. The density is the number of families.

**Table 1.** Within family coefficient of variance (CV). Size (area) measured in µm^2^, fitness as individual lambda (λ_ind_)

| Trait | Temp. (°C) | Experimental evol. regime | Within family CV (mean±SE) | Difference between evolution regimes | |
| --- | --- | --- | --- | --- | --- |
|  |  |  |  | F (ndf=1, ddf=96) | p |
| Area | 20 | Fast temp. cycles | 0.092 ± 0.009 | 3.799 | 0.054 |
| | | Increased warming | 0.115 ± 0.009 | | |
| | 25 | Fast temp. cycles | 0.100 ± 0.011 | 0.014 | 0.905 |
| | | Increased warming | 0.102 ± 0.011 | | |
| Fitness | 20 | Fast temp. cycles | 0.049 ± 0.005 | 0.266 | 0.608 |
| | | Increased warming | 0.053 ± 0.005 | | |
| | 25 | Fast temp. cycles | 0.087 ± 0.011 | 0.338 | 0.563 |
| | | Increased warming | 0.079 ± 0.011 | | |

#### Genetic variance and correlations

There was overall significant genetic variance for body size (measured as area) for the 4 combinations of temperature and evolution regime (models with genetic variance were at least 7 DIC lower compared to models without genetic variance, Table 2). There was also significant genetic covariance between temperatures for both evolution regimes (*fast temp. cycle*: model with covariance included was 2.04 DIC lower than model without covariance; *increased warming*: model with covariance was 2.79 DIC lower than model without covariance). However, the pairwise comparisons of the posterior distributions of heritability, evolvability and genetic correlations were not significantly different between the 4 different combinations of temperature and evolution regimes (all 95% credibility intervals of the pairwise differences contained zero).

**Table 2.** Heritability (H^2^), I^2^ (Evolvability, in percentage) and genetic correlations for body size (area).

| Experimental evol.<br>regime | Temp. (°C) | Posterior mode $H^2$<br>(95% CI) | Posterior mode $I^2$<br>(95% CI) | Posterior mode genetic<br>correlation across<br>temperatures (95% CI) |
| --- | --- | --- | --- | --- |
| Fast temp. cycles | 20 | 0.59 (0.44, 0.70) | 1.13 (0.56, 2.01) | 0.20 (-0.10, 0.38) |
|  | 25 | 0.56 (0.42, 0.66) | 1.30 (0.53, 2.67) |  |
| Increased warming | 20 | 0.50 (0.37, 0.65) | 1.08 (0.57, 2.42) | 0.20 (-0.12, 0.41) |
|  | 25 | 0.49 (0.42, 0.67) | 1.54 (0.69, 2.97) |  |

## Discussion

We found that evolution in an environment that changed in temperature every 2^nd^ generation (*Fast temperature cycles* regime) resulted in the evolution of increased phenotypic plasticity in body size. In contrast, we did not find any evidence of increased diversifying bet hedging in this evolutionary regime, since there was neither increased phenotypic variance within families nor reduced heritability.

Evolution in variable environments with no environmental correlations across generations is predicted to result in increased importance of either phenotypic plasticity or bet-hedging (Tufto, 2015). While phenotypic plasticity should be favored when the environment contains predictable cues for development, bet-hedging should be favored instead in less predictable environments (Botero *et al*., 2015; Tufto, 2015). Moreover, the timescale of environmental variation relative to the generation time is also important, and when modeled by Tufto, (2015), environmental changes every 2^nd^ generation is identified as the intersection between the evolution of bet-hedging, reversible plasticity or developmental plasticity. Since the *Fast temperature cycle* regime experienced fluctuations every 2^nd^ generation, they are ideally suited for investigating the evolution of plasticity and bet-hedging in adult peak body size, an irreversible plastic trait closely connected to fitness.

We found evolution of increased phenotypic plasticity in the *Fast temperature cycle regime*, manifested as a larger size difference between 20°C and 25°C (steeper reaction norm). Evolution of increased phenotypic plasticity in more variable environments is predicted by theory (Moran, 1992; Gavrilets & Scheiner, 1993; Kuijper & Hoyle, 2015), and indeed studies using natural populations (Lind & Johansson, 2007; Lind *et al*., 2011) or species (Hollander, 2008) have found a positive correlation between the degree of environmental heterogeneity and increased plasticity. Our study, using experimental evolution, supports these results and pinpoint large temporal variation between temperature extremes as the causative selection force underlying evolution of increased plasticity. This has only been showed once before, in a recent experimental evolution study of the microalgae *Thalassiosira pseudonana*, where populations evolving under fast fluctuations evolve increased plasticity in photosynthesis (Schaum *et al*., 2022). Our results also align with the recent finding that laboratory-adapted populations of Zebra fish (*Danio rerio*), evolving in very stable environments, have reduced physiological plasticity compared to their wild-caught counterparts (Morgan *et al*., 2022). Together, these studies demonstrate the importance of environmental heterogeneity for the evolution of plasticity. It should however be noted that very fast or unpredictable environmental change can make it impossible to predict the environment, and in these circumstances plasticity may be selected against (Tufto, 2015), which was recently demonstrated using experimental evolution in the microalgae *Dunaliella salina* (Leung *et al*., 2020). Increased environmental variation is however not the only factor that can influence the evolution of plasticity, but plasticity may also evolve when a population is exposed to a novel (but stable) environment (Lande, 2009; Chevin *et al*., 2010; Corl *et al*., 2018). Evolution of increased plasticity has been shown for *C. remanei* evolving in very heat-stressed environments (36.8°C), which demonstrates that evolution of plasticity also can be a way to survive novel environment (Sikkink *et al*., 2014b), a phenomenon called plasticity-led evolution (Levis & Pfennig, 2020).

In contrast, we did not find any evolution of increased diversifying bet-hedging. While empirical evidence for diversifying bet-hedging is much rarer than for phenotypic plasticity, it is also much harder to detect since it is not trait means but trait variances that needs to be measured. We have the power to detect differences in within-family CV where the selection regimes explain at least 10% of the variation, which corresponds to a large effect (Cohen, 1988). Still, there are a number of examples of bet-hedging, mainly regarding delayed germination in desert plants (Philippi, 1993; Clauss & Venable, 2000; Venable, 2007), but also diapause in fish from ephemeral pools (Furness *et al*., 2015). In addition, diversifying bet-hedging has also evolved as a result of experimental evolution in unpredictable environments in bacteria (Beaumont *et al*., 2009) and fungi (Graham *et al*., 2014). However, as predicted by Bull, (1987), a common factor in these examples is the very strong fitness differences between environments, where one environmental state (for example dry conditions) results in very low fitness. This contrasts to most examples of phenotypic plasticity, where reproduction is possible to achieve in all environments, even if they are not suitable without plastic adjustment of the phenotype. It has however been found that natural isolates of *C. elegans* differ in their variance of total reproduction, which has been suggested to result from a bet-hedging strategy where lines with more phenotypic variance may originate from more variable environments (Diaz & Viney, 2014). We did not find any difference in reproductive variance between our selection regimes, so further studies are needed to understand why natural *C. elegans* isolates differ in their phenotypic variance of reproduction.

When exposed to increasing temperatures, organisms generally show phenotypic plasticity in size, and develop faster to mature smaller. Although there are exceptions, especially in terrestrial arthropods (Horne *et al*., 2015), this relationship is so general that it is termed the temperature-size rule (Atkinson, 1994). Therefore, it is not surprising that plasticity in size was present in both evolutionary regimes, and like *C. elegans* (Kammenga *et al*., 2007), we find that *C. remanei* follows the temperature-size rule. Importantly however, the regimes differed in the degree of size plasticity.

Whether the temperature-size rule (Atkinson, 1994) reflects an adaptive or non-adaptive response to temperature is not resolved. Arguments for it being non-adaptive center around constraints related to passive by-products of other temperature dependent processes (Atkinson, 1994; van der Have & de Jong, 1996; Forster *et al*., 2011). For example, small body size can be a result of reduced growth rates which can be selected for as a by-product of increased reproductive investment in warmer temperatures (Heino & Kaitala, 1999; Walczyńska *et al*., 2015), or, alternatively, as a trade-off between growth rate and resistance to oxidative stress (Kim *et al*., 2011) which increases due to increased metabolism in higher temperatures (Birnie-Gauvin *et al*., 2017).

However, small size in warm environments can also be adaptive. For example Fryxell *et al*., (2020) showed that natural selection in warm temperatures favors smaller size in mosquitofish *Gambusia affinis* and a similar result has been reported in snails (Arendt, 2015). A possible advantage of small size in high temperatures can be a regulation of oxygen demand and supply ratio (Walczyńska *et al*., 2015). In addition, as a body composed of small cells is more efficient in oxygen diffusion (Subczynski *et al*., 1989; Verberk *et al*., 2021) there will be a particularly strong selection pressure on organisms such as *Caenorhabditis* nematodes, which have a fixed number of cells and thus the cell size determines the final body size.

To assess whether temperature-induced plasticity in size is adaptive in *C. remanei*, we compared individual fitness of different-sized individuals in both temperatures (Figure 2). We found evidence of directional selection on increased size in both temperatures, but also significant stabilizing selection within each temperature. Stabilizing selection implies that the fitness optimum in both 20°C and 25°C was present in individuals within the data size-range (as opposed to at extreme phenotypes). If small size in warm temperatures were maladaptive, we would expect the largest individuals to have the highest fitness. In contrast, we found that individuals both smaller and larger than the optimum size had decreased fitness. This optimum size in the warm temperature was also substantially smaller than the optimum size at the normal temperature, thus the plastic response to decrease size as a response to warm temperature must be considered adaptive in *C. remanei*.

Interestingly, because most individuals raised in 25°C exhibit smaller size than would be optimal (Figure 2.; mean size is smaller than optimal size), we can consider this temperature plasticity to be a hyperplastic response, a special case of plasticity when plastic response overshoots the optimum and brings individuals to the other side of the new fitness peak (King & Hadfield, 2019). Since plasticity nevertheless increases fitness (compared to a hypothetical non-plastic genotype), this hyperplasticity should still be considered adaptive. In addition, we also found linear selection for large size in 20°C with individuals raised in 20°C also having slightly smaller size than would be optimal. A possible explanation is a sexual conflict between male and female worms, as males’ optimal size is smaller than females’ optimal size in *C. remanei* (Stångberg *et al*., 2020) so that males drag females from their phenotypic optimum.

In contrast to size, we didn’t find any difference in individual fitness between the regimes. While warm temperature caused a drop in total reproduction in both regimes, individuals raised in 25°C had significantly higher rate-sensitive individual fitness, which is a consequence of the temperature-induced alteration of the reproductive schedule, including a faster development time (Sekajova *et al*., 2022).

Previous experimental evolution and artificial selection studies in the SP8 line of *C. remanei*, which was our founder population, have documented fast evolutionary responses to selection in life history traits, suggesting a large amount of standing genetic variation (Chen & Maklakov, 2012; Zwoinska *et al*., 2016; Lind *et al*., 2020). Accordingly, we found substantial genetic variation for size, for all treatment × temperature combinations, which not only allowed the lines to respond to selection, but also represents a potential for further evolution. Since we used full-sibs, our estimates of genetic variance could potentially be inflated by the presence of dominance variance and epistatic interactions. Epistatic interactions are present for body size in the sister species *C. elegans* (Noble *et al*., 2017; Maulana *et al*., 2022), but while most genetic variance for this trait was found to be additive and with similar additive heritability to our estimate using an experimental evolution design (Noble *et al*., 2017), assessments of narrow-sense heritability using recombinant inbreed lines found that additive effects played a smaller role for body-size (Maulana *et al*., 2022). Therefore, we assume that our broad sense heritability overestimates the additive genetic effect, but to an unknown degree. Moreover, we didn’t observe any significant differences in broad-sense heritability between the lines, which further support our evidence of no evolution of diversified bet-hedging, which comes with the prediction of lowered heritability in traits (Tufto, 2015), nor did we observe any genetic correlations between trait values in the two temperatures, suggesting that traits can largely evolve independently in each environment.

## Conclusion

To summarize, we found that 30 generations of experimental evolution in a heterogeneous environment (*Fast temperature cycles)* resulted in the evolution of increased phenotypic plasticity, compared to evolution in a slowly changing environment (*Increased warming)*. We showed that plasticity followed the temperature size rule and was adaptive. In addition, substantial amount of standing genetic variation found in the line represents a potential for further evolutionary change.

## Author contributions

ZS, MIL, IIR and EB designed the experiment, ZS, ER and MT-B performed the phenotypic assays with the aid of MIL. EIFF and ZS analysed the data, with the aid of EB. ZS, MIL and EIFF drafted the manuscript. All authors contributed to the revision of the manuscript.

## Acknowledgements

We thank Alexei Maklakov for access to the experimental evolution lines. This work was financed by the Swedish Research Council (Vetenskapsrådet, grant no. 2016-05195 and 2020-04388) and the Carl Tryggers Foundation (grant no. CTS 17:285) to MIL.

## Data Availability Statement

Upon acceptance, the data will be archived at *Figshare*

