## Supplementary materials for "Evolution of phenotypic plasticity during environmental fluctuations"

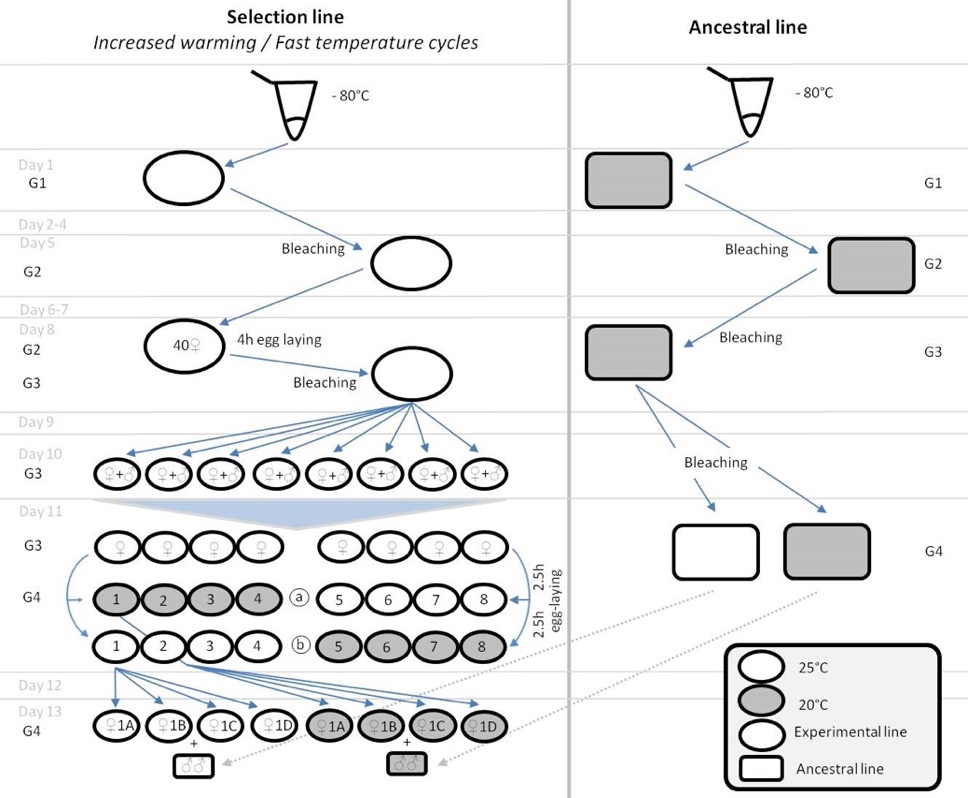

**Supplementary figure 1**: Experimental set up

Selection Line

G1: The worms are revived from freezing and placed into 25°C. After reaching sexual maturity, they are synchronized by bleaching which kills the worms with only eggs surviving.

G2: After the eggs hatch and reach their peak reproduction (day 3 after bleaching), 80 females are picked on random and transferred to a new set of 92 mm plates so that each plate contains 40 females. After 4 hours of a synchronized egg lying, the plates are treated by bleaching resulting in age-synchronized generation 3.

G3: 42 hours after the bleaching 8 virgin males and 8 virgin females are transferred to 35 mm petri plates (each plate containing one female and one male). After 21 hours, all the 8 females are mated and transferred to a new set of plates to lay eggs. Egg lying worms are kept on 35 mm plates (a) for 2.5 hours and then transferred to the new set of plates (b) for another 2.5 hours after which adult females are removed.

G4: From each female we thus obtained two sets of plates (a and b, 16 plates in total) containing approximately 20 eggs, (full siblings). The plates are moved to either 20°C or 25°C climate chamber so that for the half of the females, the set of the plates a is moved to 20°C and the set of plates b to 25°C and opposite for the other half. After another 41 hours (in 25°C) and 54 hours (in 20°C) 4 virgin females from each plate are selected on random and transferred to a new set of the plates together with 2 males from ancestral line. This results in 8 plates per family (originating from the same mother), half being in 20°C and half in 25°C which makes 64 plates (32 plates in each temperature) in total per each line and replicate.

Ancestral line

G1: The worms are revived from freezing and placed into 20°C.

G1-G3: the worms experience three cycles of bleaching which corresponds to 1 cycle of bleaching and 2 cycles of the synchronized egg laying of the selection lines.

G4: After the bleaching, the worms are split into 20°C and 25°C to be mated with females from the selection regimes.

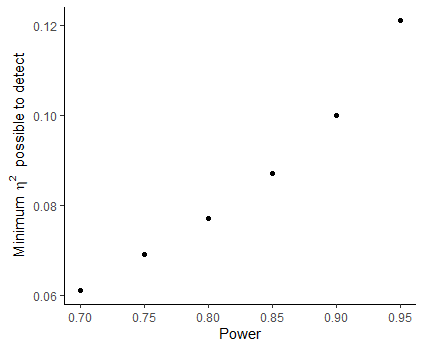

**Supplementary figure 2.** Minimum effect size (η^2^) possible to detect in ANOVA across power, given a sample size of N = 48 for each of the two selection regimes and a significance level of 0.05.
